# Transposons modulate transcriptomic and phenotypic variation via the formation of circular RNAs in maize

**DOI:** 10.1101/100578

**Authors:** Lu Chen, Pei Zhang, Yuan Fan, Juan Huang, Qiong Lu, Qing Li, Jianbing Yan, Gary J. Muehlbauer, Patrick S. Schnable, Mingqiu Dai, Lin Li

**Affiliations:** National Key Laboratory of Crop Genetic Improvement, Huazhong Agricultural University, Wuhan 430070, China.; Department of Agronomy and Plant Genetics, University of Minnesota, Saint Paul, MN 55108 USA.; Department of Plant and Microbial Biology, University of Minnesota, Saint Paul, MN 55108 USA.; Department of Agronomy, Iowa State University, Ames, Iowa 50011, USA

**Keywords:** Transposons, LINE1, Circular RNAs, Phenotypic Variation, Maize

## Abstract

Circular RNAs (circRNAs) are covalently closed, single-stranded RNA molecules. Recent studies in human showed that circRNAs can arise via transcription of reverse complementary pairs of transposons. Given the prevalence of transposons in the maize genome and dramatic genomic variation driven by transposons, we hypothesize that transposons in maize may be involved in the formation of circRNAs and further modulate phenotypic variation. To test our hypothesis, we performed circRNA-Seq on B73 seedling leaves and integrate these data with 977 publicly available mRNA-Seq datasets. We uncovered 1,551 high-confidence maize circRNAs, which show distinct genomic features as compared to linear transcripts. Comprehensive analyses demonstrated that LINE1-like elements (LLE) and their Reverse Complementary Pairs (LLERCPs) are significantly enriched in the flanking regions of circRNAs. Interestingly, the accumulation of circRNA transcripts increases, while the accumulation of linear transcripts decreases as the number of LLERCPs increases. Furthermore, genes with LLERCP-mediated circRNAs are enriched among loci that are associated with phenotypic variation. These results suggest that LLERCPs can modulate phenotypic variation by the formation of circRNAs. As a proof of concept, we showed that the presence/absence variation of LLERCPs could result in expression variation of one cicrRNA, circ352, and further related to plant height through the interaction between circRNA and functional linear transcript. Our first glimpse of circRNAs uncovers a new role for transposons in the modulation of transcriptomic and phenotypic variation via the formation of circRNAs.

## Introduction

RNAs were primarily defined by the central dogma as messenger molecules in the process of gene expression. However, ample evidence suggests that RNAs function not only in protein bisosynthesis in a coding manner, but also as regulatory molecules in a non-coding manner, which enhances the importance of RNA species (Morris and Mattrick 2014; Rinn and Chang 2012). There are many types of non-coding RNAs, including microRNAs, small interfering RNAs, natural cis-acting siRNAs and other long noncoding RNAs. One rising star of these non-coding RNA species is circular RNAs (circRNAs) (Chen et al 2015). CircRNAs are covalently closed, single-stranded RNA molecules with a 3’, 5’ – phosphodiester or a 2’, 5’ – phosphodiester bond at the junction site (Chen and Yang 2015). Approximately 30 years ago a handful of circRNAs were identified and thought to be transcriptional noise due to aberrant splicing and lack of identified functional role (Hsu and Coca-prados, 1979; Nigro et al. 1991; Capel et al. 1993; Zaphiropoulos 1993; Pasman et al. 1996). Recently, circRNAs have been reported to influence the expression of parental linear transcripts and other genes (Chen et al. 2015).

With the advent of genomic sequencing techniques and high-efficiency bioinformatic analysis, genome-wide scans for circRNAs in different cell types or tissues have been conducted in a wide range of species (Chen et al. 2015). Although most circRNAs were expressed at low levels, thousands of circRNAs have been identified in mouse, *Caenorhabditis elegans,* Drosophila, and humans (Salzman et al. 2012; Memczak et al. 2013; Ashwal et al. 2014; Westholm et al. 2014; Guo et al. 2014; Bachmayr-Heyda et al. 2015; Gao et al. 2015) and dozens of circRNAs were shown to be highly expressed in a cell type- or tissue-specific manner, suggestive of functionality. CircRNAs have been identified across the eukaryotic tree of life, indicative of evolutionary conserved feature of eukaryotic gene expression process (Wang et al. 2014). More evidence has been accumulated to indicate that circRNAs have the potential to shape gene expression as miRNA sponges, regulating transcription and interfering with splicing (Kulcheski 2016). More recent mammalian studies have shown that transposons are enriched in the flanking regions of circRNAs and have the potential to mediate the formation of circRNAs via reverse complementary pairs (Jeck et al, 2013; Zhang et al, 2014).

Recently, identification and characterization of plant circRNAs has been conducted. Kulcheski and colleagues (2014) performed the first genome-wide scan in *Arabidopsis* and identified three circRNAs. In rice, 2,354 circRNAs were identified via deep sequencing and computational analysis of strand-specific RNA-Seq data (Lu et al., 2015). Meanwhile, Ye and colleagues (2015) performed a comparative transcriptomic analysis of circRNAs, identified 12,037 and 6,012 circRNAs in rice and *Arabidopsis*, respectively, indicative of widespread occurrence of circRNAs in plants. Plant circRNAs were shown to be conserved, expressed at low levels and in a tissue-specific manner, and were found to be differentially expressed under phosphate-sufficient and starvation conditions (Ye et al. 2015). In contrast to what has been observed in mammals, repetitive elements and reverse complementary sequences are not significantly enriched in the flanking regions of circRNAs in rice and *Arabidopsis* (Lu et al. 2015; Ye et al. 2015).

Maize is one of the most widely grown crops in the world. Although the first version of maize reference genome was released in 2009, the genome-wide annotation of functional elements is still ongoing (Schnable et al. 2009). For example, Li and colleagues (2014) identified and characterized over 20,000 long noncoding RNAs (Li et al. 2014). Increased complexity of the maize transcriptome has been revealed through third generation sequencing (Wang et al. 2016). Until recently, transcriptomic studies were focused on genes with linear transcripts (both protein coding and noncoding), ignoring the widely existing noncoding circRNAs in maize. Meanwhile, more than 85% of the maize genome is repetitive transposable elements (Schnable et al. 2009). Transposons have been reported to be involved in the regulation of functional gene expression and phenotypic variation by either insertion in coding, regulatory and intergenic regions or alternation of epigenomic variation, or large scale genomic transposition (Lisch 2009; Lisch 2013; Wei and Cao 2016). A recent study showed that human reverse complementary pairs of transposons can mediate the formation of circRNAs (Zhang et al. 2014). Given the prevalence of transposons in the maize genome (Schnable et al., 2009) and pilot discovery of circular DNA elements mediated by Mu transpositions in maize (Sundaresan and Freeling 1987), we would expect ample circRNAs in maize. Additionally, maize transposons vary dramatically among haplotypes (Fu et al. 2002; Bennetzen 2005; Dooner et al. 2008), which may lead to the extensive diversity of circRNAs among haploypes. Together with the evidence that circRNAs can function as miRNA sponges, and transcriptional regulators of their parental functional linear transcript isoforms (Chen et al. 2015), we predict that transposons could function in a new manner to modulate phenotypic variation via the formation of circRNAs.

Maize is a widely studied model for genetics. Diverse mRNA-Seq datasets have been deposited in the short read archive database. Although it is thought that circRNAs lack poly(A) tails, some circRNAs have also been detected in poly(A)-selected libraries (Salzman et al. 2012). To identify circRNAs, circRNA-Seq, which combines RNase R treatment and Next Generation Sequencing (NGS), have been developed (Jeck et al. 2013). Here, we collected a comprehensive transcriptome dataset and identified 1,551 high-confidence circRNAs in maize, which showed distinct genomic features around the junction loci. Interestingly, unlike rice and *Arabidopsis,* retrotransposon LINE1-like elements (LLEs) and their reverse complementary pairs (LLERCPs) are likely to affect the formation of circRNAs. The dramatic genomic variation of LLEs among diverse inbreds is significantly associated with the expression-level variation of circRNAs and parental linear RNAs, which is further related to agronomic trait variation. Our study provided a first glimpse of maize circular RNAs, and uncovered a novel functional role of transposons in the regulation of transcriptomic and phenotypic variation in maize probably via the formation of circRNAs.

## Results

### Circular RNAs exist widely in maize

CircRNAs can be classified into intergenic and genic groups according to their origin. In this report we mainly focus on genic circRNAs, because they were more likely to be directly associated with gene expression and phenotypic variation. To detect circRNAs, we performed circRNA-Seq on 2-week old seedlings of the maize inbred line B73. We also collected a comprehensive transcriptome dataset, including 977 publicly available RNA-Seq datasets derived from diverse B73 tissues and from a diverse panel of maize inbreds (Hirsch et al. 2014; Fu et al. 2014). A series of circRNA identification and quality-control steps were implemented to ensure only the inclusion of high-confidence circRNAs for further analysis (Fig S1; see Methods). These analyses revealed two major types of genic circRNAs – exonic circRNAs and intronic circRNAs (Fig 1A). Totally, we uncovered 1,551 species of circRNAs, of which 1,453 are of exonic origin and 98 are intronic circRNAs (Fig 1D; Table S1). These high-confidence circRNAs, which have at least 2 reads covering the backsplicing junction, were distributed across all the whole genome, however, similar to genes, more circRNAs were from both ends of chromosomes (Fig S2).

**Fig 1.**
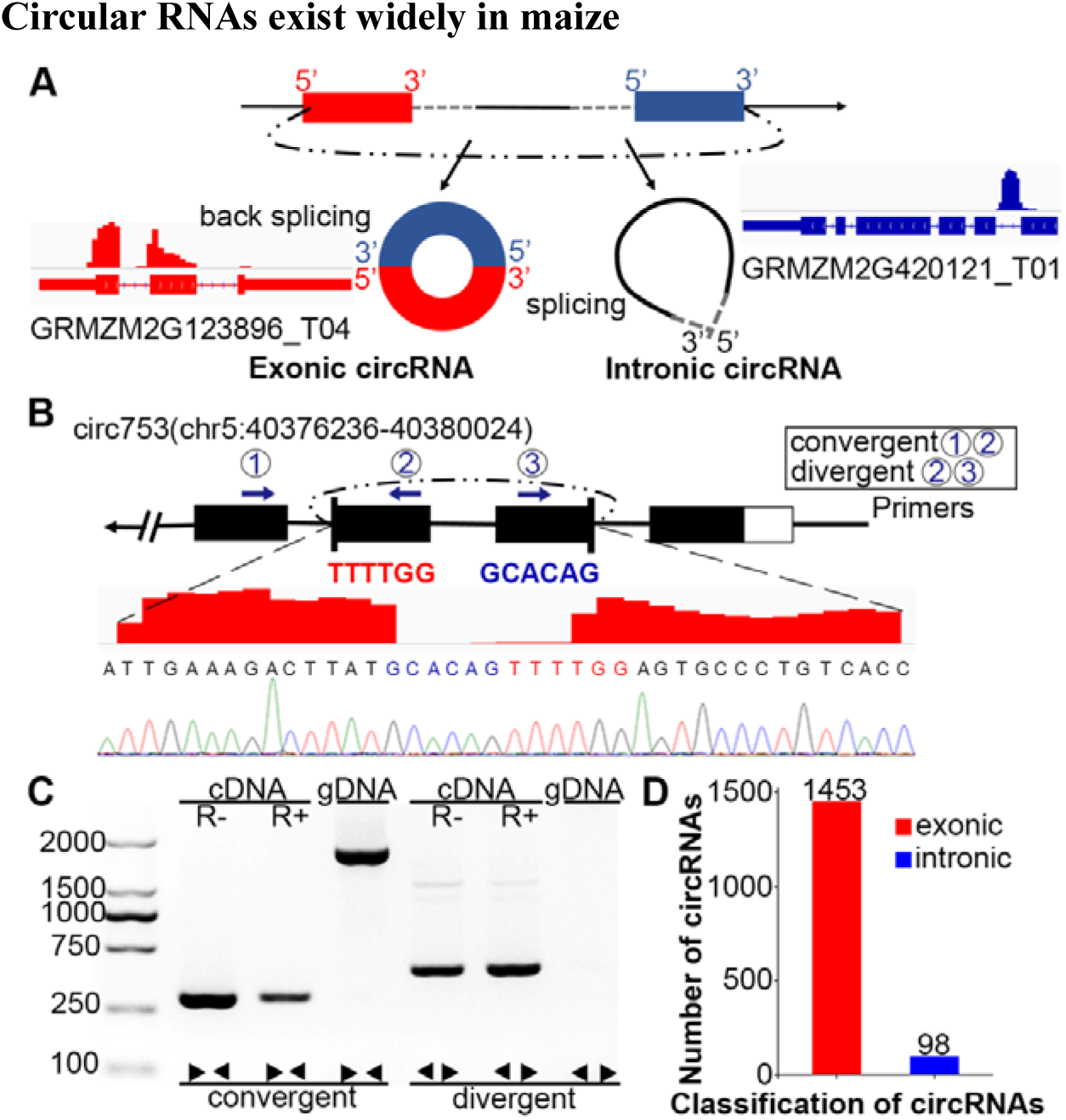
Genome-wide identification of circRNAs in maize. (A) Origins of exonic and intronic circRNAs. The black dashed line indicates backsplicing junction site, which also represents the circRNA junction site. The grey dashed line and the black line designate an intron. Red and blue rectangle represent exons. (B) (C) An example of circRNA that was validated via amplification and sequencing. R+, R- represent samples with and without RNase R treatment, respectively. Arrows (①, ②, ③) designate PCR primers. (D) Number of circRNAs identified.

In an attempt to validate high-confidence circRNAs, we amplified B73 leaf cDNAs from total RNAs or from RNAs of which the linear mRNAs have been removed by RNase R treatment, as well as genomic DNA using pairs of divergent and convergent primers for 19 randomly-selected circRNAs (Fig 1B). All convergent primers successfully amplified transcribed fragments with the expected length. Meanwhile, all 19 pairs of divergent primers yielded amplification products from both cDNAs that had and had not been treated with RNase R but not from the genomic DNAs. All the amplification products from the divergent primers were shown to be derived from the target regions of circRNAs (i.e., spanning the junction sites) via Sanger sequencing (Fig 1C; Fig S3; Table S2; see Methods). These results establish the reliability of our high-confidence circRNA discovery pipeline.

### Circular RNAs and their parental gene loci show distinct characteristics as compared to linear genes

To characterize circRNAs and their genomic loci, we compared circRNAs with the linear transcripts from which they were derived and randomly-selected genes without detectable circRNAs. Similar to previous reports, maize circRNAs exhibit distinct features. Genes with detectable circRNAs are significantly longer than randomly-selected genes (Fig 2A). Overall, the average exon length of circRNAs is significantly shorter than that of randomly-selected linear transcripts, especially for circRNAs with multi exons. However, for circRNAs with only a single exon, the exon length is significantly longer than that of randomly-selected linear transcripts (Fig 2B). Interestingly, the exon length of circRNAs decreases along with the increase of exon number when exon number of circRNAs is less than five, then enters a plateau (Fig 2C). Notably, the flanking intron length of the junctions of circRNAs is significantly larger than that of linear transcripts of randomly-selected gene loci (Fig 2D). Although circRNAs themselves are likely to be expressed at low levels, most of their parental genes are likely to be highly expressed (Fig 2E). Notably, there is no obvious correlation between the accumulation of circRNAs and their corresponding parental genes (Fig 2F), suggesting complicated relationships of these two RNA types in maize. Additionally, a significantly higher proportion of circRNAs could be aligned with miRNAs than that of the genome-wide linear transcripts, indicative of potential miRNA mimic capability (152/1245) (Fig S4). Furthermore, circRNAs are more likely to be expressed in a tissue-specific manner than are genome-wide linear transcripts (Fig S5). All these features of maize circRNAs are similar to that described in other species, suggesting that fundamental features and regulation of circRNAs exhibit evolutionary conservation.

**Fig 2.**
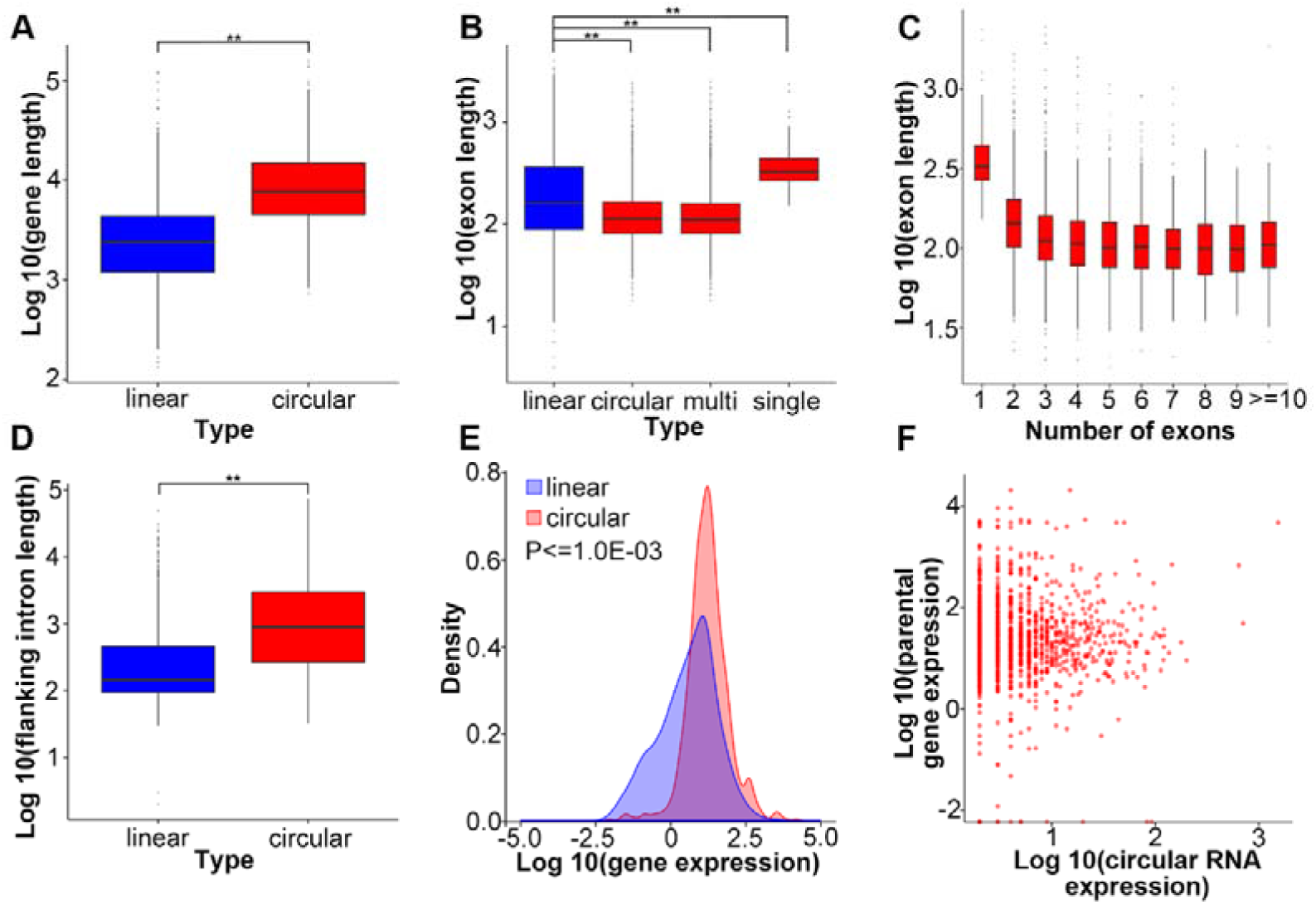
Distinct genomic features of circRNAs in maize. (A) Comparison of gene lengths between genes with and without detected circRNAs. Linear and circular represent randomly-selected genes without detectable circRNAs and genes with detectable circRNAs in B73 v3.26 annotation, respectively. (B) Comparison of exon length between exons with and without detectable circRNAs. Linear indicated random selected exons without detectable circRNAs, circular indicates exons with detectable circRNAs, while multi and single represent circRNAs with multiple exons and single exon, respectively. (C) Length distribution of exons of circRNAs. (D) Length distribution of flanking introns of genes with and without detectable circRNAs. Y-axis in (B), (C) and (D) indicates the log 10 of exon or intron length (bp). (E) Comparison of expression levels (FPKM) between genes with and without detectable circRNAs. Linear and circular represent randomly-selected genes without detectable circRNAs and genes with detectable circRNAs in B73 v3.26 annotation, respectively. (F) Expression-level relationships between circRNAs (junction reads) and their parental genes (TPM).

### Retrotransposon LINE1-like elements (LLEs) and their reverse complementary pairs (LLERCPs) are significantly enriched in the flanking regions of circRNAs

To test if transposons play a role in the formation and expression of circRNAs, we annotated 35 kb of genomic sequences upstream and downstream of the circularization junctions (Fig 3A) for repetitive elements using RepeatMasker (see Methods). Interestingly, retrotransposon LINE1-like elements (LLE) were significantly enriched in these flanking genomic regions, spreading up to 25 kb upstream and 30kb downstream of the circRNA junctions (Fig 3A). Like Alu sequences in humans (Zhang et al., 2014), we also observed divergent and convergent distribution pattern of LLEs around circRNAs, ensuring the formation of reverse complementary pairs of LLEs (LLERCPs). LLERCPs may form a stem-loop structure, of which the stem can be recognized and excised for the formation of circRNAs (Fig 3B). In addition, LLERCPs exhibit greater enrichment upstream and downstream of circRNAs than do LLEs *per se* (4.5-fold change *vs.* 20% more; Fig 3C, D), consistent with a role for LLERCPs in the process of RNA circularization (Fig 3B).

**Fig 3.**
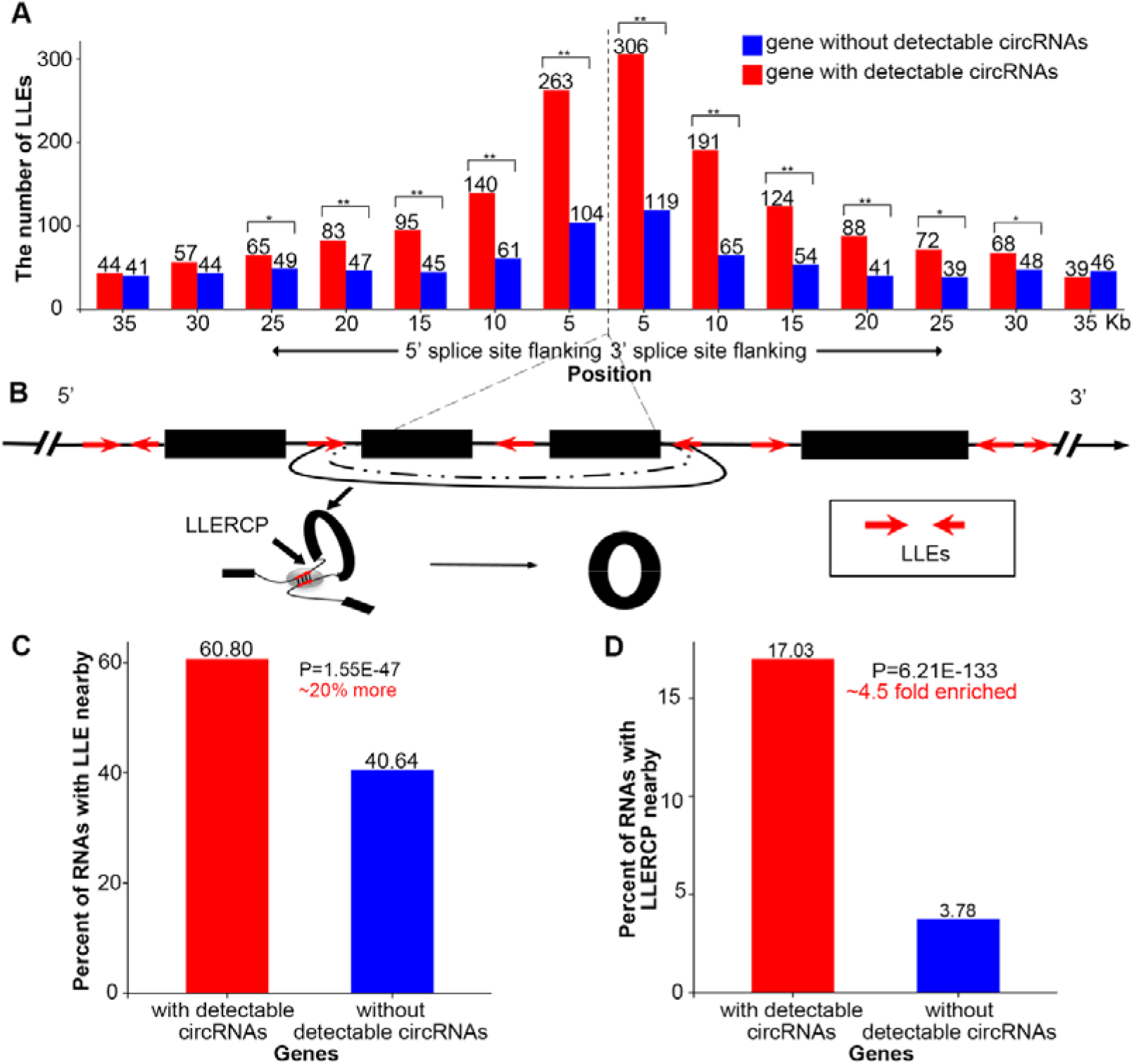
Enrichment of retrotransposon LINE1-like elements (LLEs) and their reverse complementary pairs (LLERCPs) around circRNAs. (A) Number of LLEs in the flanking regions of circular RNAs and randomly-selected linear RNAs without detectable circRNAs. Two asterisks represent a P value less than 0.01, one asterisk represents P value less than 0.05, according to the 1000 time simulations. (B) Schematic diagrams showing LLERCPs and formation of circRNAs. LLERCPs may form stem-loop structure during the formation of circRNAs. The solid black line linking two red arrows indicates that two LLEs have a reverse complementary relationship. The dashed black line indicates the back-splicing site of the circRNA. The gray oval in the stem-loop structure designates the reverse complementary region of LLERCPs. The back arrow in the stem-loop structure shows the back-splicing process required for formation of circRNAs. (C) (D) Enrichments of LLEs and LLERCPs in the flanking regions of genes with or without circRNAs.

### Reverse complementary pairs of LINE1-like elements (LLERCPs) are associated with the expression of circular and linear RNAs

To determine if LLERCPs affect the accumulation of circRNAs, we identified 27 genes for which both circRNAs with LLERCPs and circRNAs without LLERCPs have been detected (Fig 4A). A paired T-test for these 27 genes showed that circRNAs with LLERCPs accumulated to higher levels than did circRNAs without LLERCPs nearby (35kb), indicating that LLERCPs could reinforce the expression of circRNAs (Fig 4B). Most interestingly, although the expression-level of different genes inherently varies, along with the increase of the number of LLERCPs, the expression levels of corresponding circRNAs increase, confirming the incremental role of LLERCPs in circRNAs (Fig 4C). In contrast to circRNAs, the expression-levels of linear transcripts decrease as the number of LLERCPs increase (Fig 4D). The complicated pattern of expression-level variation for circRNAs and linear transcripts along with the increase of LLERCPs may indicate the complex interaction between circRNAs and linear RNAs. Taken together, these results indicate that the number of LLERCPs may play a role in the regulation of the expression of circRNAs and linear RNAs.

**Fig 4.**
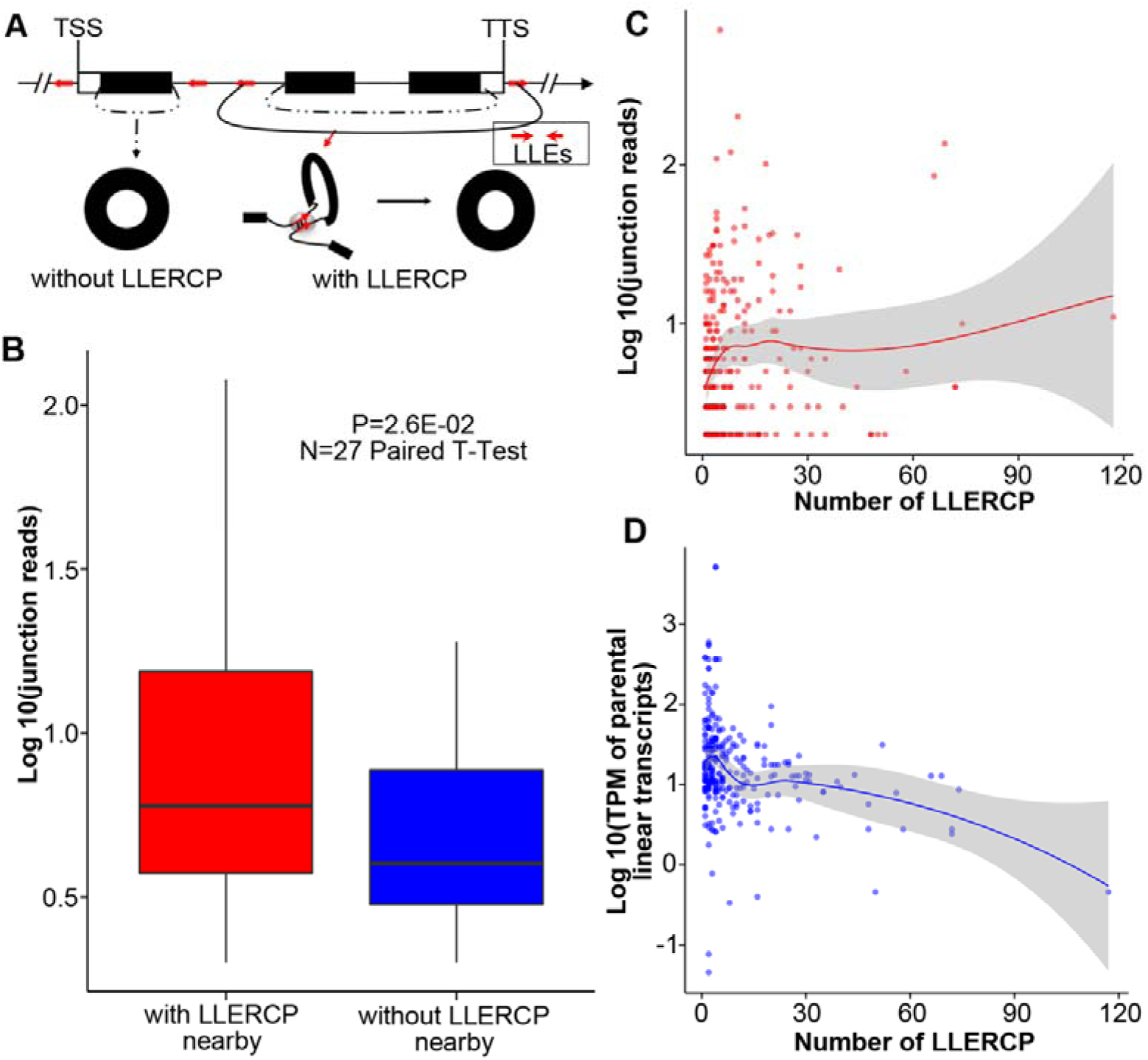
The number of Reverse complementary pairs of LINE1-like elements (RCPLLEs) is associated with the expression of circRNAs and linear transcripts. (A) A schematic diagram of the origination of circRNAs. There are some loci, where both circRNAs without intact LLERCPs and circRNAs with LLERCPs are observed. LLERCPs may form stem-loop structures prior to recruitment of splicesomes during the formation of circRNAs. (B) Expression levels of circRNAs with LLERCPs nearby are significantly higher than those of circRNAs without intact LLERCPs nearby. (C), (D) Expression-level variation of circRNAs and their parental linear transcripts along with the increase of RCPLLEs. Red and blue dotted lines link the average value of each group. The number under each violin plot shows the number of circRNAs (C) or parental linear transcripts (D) in each RCP group.

### Reverse complementary pairs of LINE1-like elements are less likely to be associated with small RNAs in maize

LLERCPs may form stem-loop structures during the formation of circRNAs (Fig 5A). These stem-loop structures are similar to the hairpin structures associated with the formation of small RNAs. To test the relationship between small RNAs and LLERCPs, we downloaded the small RNA-Seq dataset from maize B73 5d shoots, mapped small RNAs against the whole maize genome, and quantified the accumulation levels of 21nt to 25nt species of small RNAs within all LLEs (Regulski et al. 2013). Overall, the normalized number of small RNA reads that aligned to genic RCP LLEs is less than the number that aligned to genic nonRCP LLEs (Fig 5B; 5C). Specifically, the accumulation levels of 24nt small RNAs on genic RCP LLEs are significantly lower than that of randomly-selected genic nonRCP LLEs. These results indicate that LLERCPs may provide a more stable condition of precursor transcripts for the subsequent formation of circRNAs.

**Fig 5.**
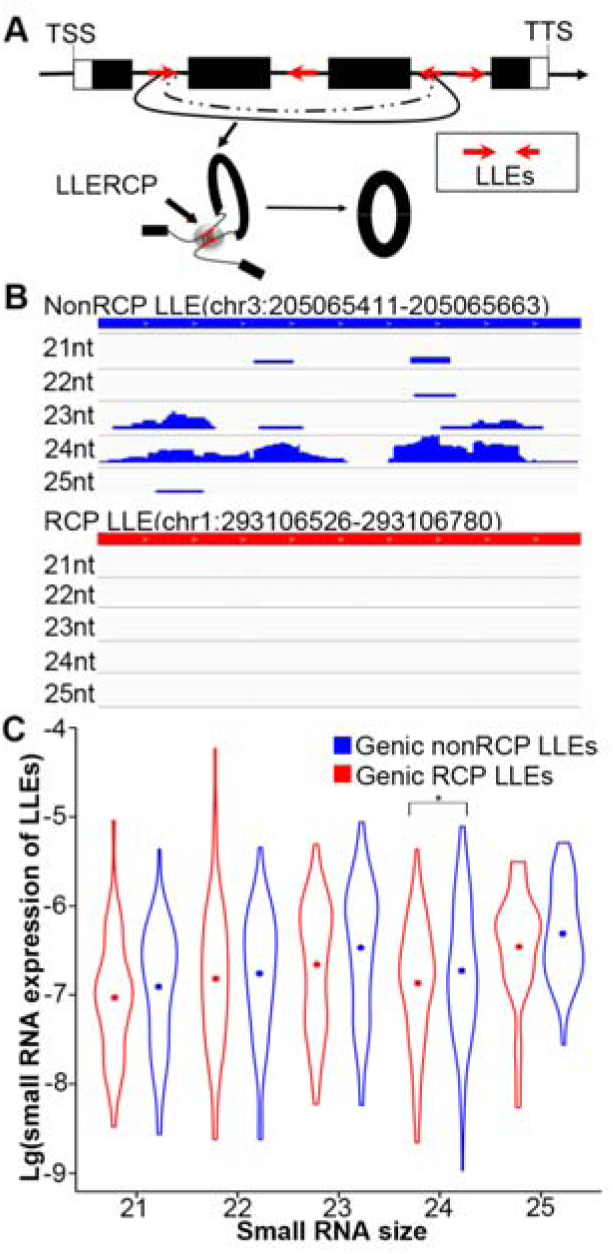
Depletion of small RNAs from RCP LLEs. (A) A schematic diagram of LLERCPs that might be involved in the formation of circRNAs in maize. LLERCPs might form a stem-loop structure, which may either be excised for the biogenesis of small RNAs, or provide a stable structure to recruit the splicesome for formation of circRNAs. (B) An example of alignment of small RNAs in genic RCP LLEs and randomly-selected genic non-RCP LLEs. Randomly-selected genic non-RCP LLEs had more small RNAs aligned than genic RCP LLE, indicative of depletion of small RNAs (especially 24 nt species) in RCP LLEs. (C) Normalized accumulation of genic RCP LLE-derived 24nt small RNAs is significantly higher than that of genic non-RCP-derived small RNAs. The asterisk represents the significant level less than 0.05.

### Variation in LLERCP content is associated with phenotypic variation

A series of analyses were conducted to test for potential functional roles of circRNAs. First, a gene ontology enrichment analysis showed that genes that give rise to circRNAs are more likely to be associated with cellular component organization, intracellular membrane-bounded organelle, nucleotide binding, helicase activity and coenzyme binding than the reference genes in B73 genome (Fig S6). Additionally, as compared to genes without detectable circRNAs, genes with detectable circRNAs were enriched to overlap a set of 6,575 non-redundant genes that have been associated with phenotypic variation based on genome-wide association mapping (GWAS) (see Methods), as tested via Monte Carlo simulation (P<=0.001; see Methods). The proportion of genes with circRNAs and functional association sites is significantly higher than the randomly-selected genes without detectable circRNAs (P = 5.22E-31), randomly-selected genes with similar gene length distribution as circRNA genes (P = 1.09E-03), and randomly-selected genes with LLERCPs but without detectable circRNAs (P=3.34E-14; Fig 6A). More intriguingly, about 17% of genes with circRNAs and GWAS signals are the ones with LLERCPs (Fig 6B). Such an enrichment of LLERCPs in genes with circRNAs and GWAS signals is much higher than the randomly-selected genes without detectable circRNAs (P = 6.45E-26). Meanwhile, such an enrichment is also observed compared to genes without detectable circRNA but with similar gene length distribution as circRNA genes (P = 6.98E-06; Fig 6B). Additionally, circRNAs with miRNA targets is also more likely to be associated with phenotypic variation (Fig S7).

**Fig 6.**
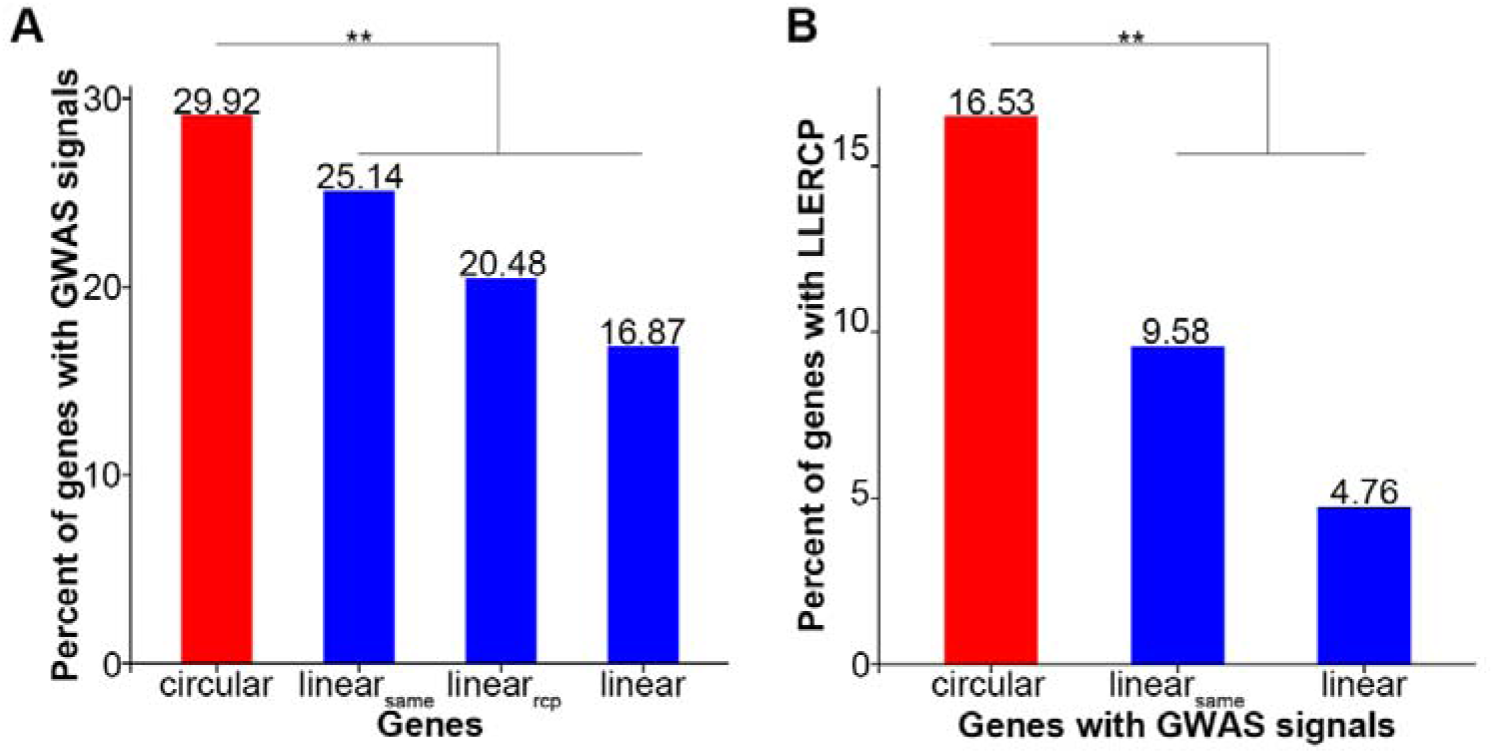
Genes associated with phenotypic variation are enriched among genes with detectable circRNAs and LLERCPs. (A) Enrichment of GWAS signals in genes with detectable circRNAs. (B) Enrichment of LLERCPs in genes with detectable circRNAs and GWAS signals. Circular, linear_same_, linear_rcp_ and linear represent genes with detectable circRNAs, randomly-selected genes with same gene-length distribution as circRNA genes, randomly-selected genes with LLERCPs but without detectable circRNAs, and randomly-selected genes, respectively. The asterisks represent the significant level less than 0.01.

If circRNAs contribute to phenotypic variation we would expect that LLERCPs would exhibit presence/absence polymorphism in the trait-associated genes among the lines used in the GWAS and that this variation would itself be associated to phenotypic variation. To test this hypothesis we profiled a gene, *GRMZM2G089149* (Fig 7A), that was significantly associated with plant height in a US association panel (Wallace et al. 2013) and that contains LLERCPs and accumulates a circRNA, circ352, for variation in LLERCPs among 38 diverse inbreds in a Chinese association panel (Fig 7B). This plant-height associated gene, *GRMZM2G089149,* has 13 exons and 12 introns, of which the 3^rd^ and 11^th^ are the biggest introns. CircRNA, circ352, spans exons from 4^th^ to 11^th^, and is associated with the LLERCPs within 3^rd^ and 11^th^ introns. Genomic amplification and quantitative real-time PCR analysis showed that all inbreds having LLERCPs in *GRMZM2G089149* had detectable levels of circ352, while only 10% of the inbred lines that lacked LLERCPs accumulate detectable levels of circ352 (Fig 7C). These results provide evidence that presence/absence of LLERCPs is related to the formation of circRNAs. Further, the expression of circ352 is positively associated with the level of the linear transcript of *GRMZM2G089149* (Fig 7D), suggesting that the expression of circRNAs may have functional consequence on the linear transcripts. More interestingly, inbreds with detectable LLERCPs are significantly shorter than those without detectable LLERCPs (P = 4.0E-02; Fig 7E), consistent with the hypothesis that the presence/absence of LLERCPs affects variation in plant height. We did not, however, observe a correlation between plant height and the accumulation of linear or circular transcripts across inbreds (Table S3). This might be due to that the tissue analyzed (topmost leaf of 2-week seedlings) was not associated with plant height. Even so, these results provide intriguing clues about a mechanism by which transposons could play a functional role in the phenotypic variation via the formation of circRNAs.

**Fig 7.**
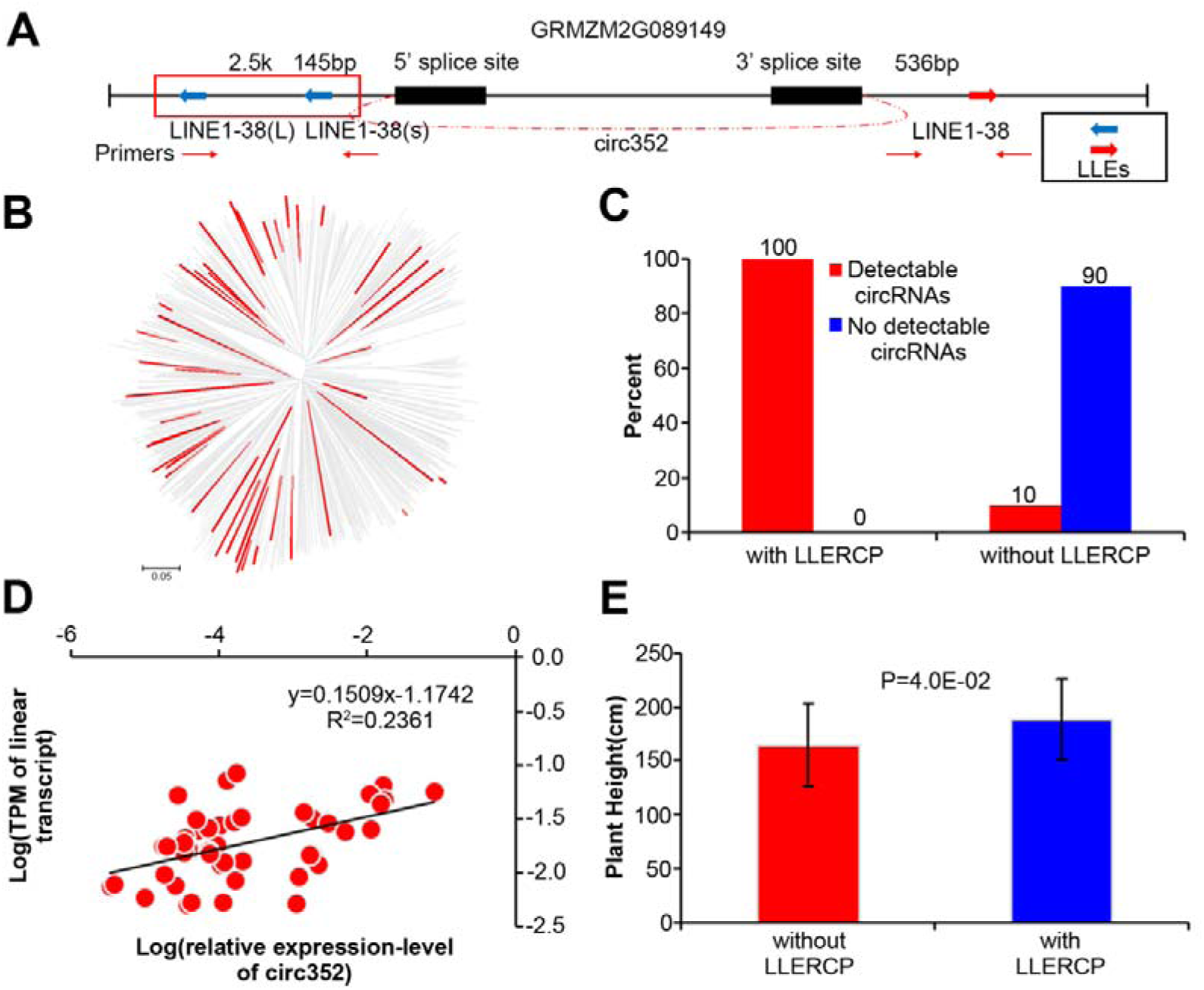
The presence/absence of RCPLLE is associated with variation in plant height among inbreds. (A) circ352 is derived from the gene *GRMZM2G089149.* The B73 allele of *GRMZM2G089149* contains an intact RCPLLE. The red box indicates that two LLEs were amplified by one pair of primers together. (B) Thirty-eight diverse inbreds (in red) were randomly-selected for analysis from a larger association mapping panel in maize (Yang et al. 2011). (C) The presence/absence of RCPLLEs in *GRMZM2G089149* is associated with the accumulation of detectable levels of circ352 among inbreds. (D) The accumulation of circ352 is associated with that of the linear transcript from *GRMZM2G089149.* (E) Inbreds with LLERCP are significantly shorter than those that do not contain RCPLLEs.

## Discussion

### Widespread existence of circRNAs in Eukaryotes

With the advent of next generation sequencing, ENCODE Projects of humans and mouse have been completed, which have dramatically revolutionized our understanding of eukaryotic transcriptome (Djebali et al. 2012; Dunham et al. 2012; Yue et al. 2014). Many lines of evidence have shown the prevalence of transcription from the whole genome, leading to the discovery of hundreds of thousands of long noncoding RNAs (lncRNAs). LncRNAs have been shown to function as super regulators at the genome, transcriptome, protein, and metabolic levels (Rinn and Chang 2012). As one type of the lncRNAs, circRNAs were identified over 30 years ago (Hsu and Coca-prados, 1979). However, due to the limitation of molecular techniques and the structure of non 3’ poly(A) tail and 5’ end caps, only a handful of circRNAs have been identified and were considered as transcriptional noise (Hsu and Coca-prados, 1979; Nigro et al. 1991; Capel et al. 1993; Zaphiropoulos 1993; Pasman et al. 1996). Recently, with the development of circRNA-Seq, thousands of circRNAs have been identified in animals and plants (Jeck et al. 2013; Chen 2016). These circRNAs are much more stable than linear transcripts, conserved among species, and their flanking sequences are associated with Alu elements in animals (Jeck et al. 2013). In Rice and Arabidopsis, over 2,000 circRNAs have been identified, showed distinct genomic features compared to linear genes and transcripts (Ye et al. 2015). In maize, we collected a comprehensive dataset from different Next Generation Sequencing (NGS) techniques, including traditional linear mRNA-Seq and circRNA-Seq with circRNA enrichments, and identified over 2,000 circRNAs in maize, of which 1,551 were high-confidence circRNAs. Taken together, detection of circRNAs in mammals and plants suggests that circRNAs are ubiquitous, and thus are not likely transcriptional noise or a by-product of transcription of functional linear transcripts, but function as miRNA mimics and regulators of gene expression (Chen 2016).

Our first glimpse of maize circRNAs not only detected thousands of circRNAs, but also found many interesting aspects of circRNAs in maize. Maize circRNAs show distinct features, which are in agreement with that of circRNAs in other species. Most strikingly, although transposons and reverse complementary pairs of repetitive elements have not shown to be enriched in the flanking regions of the circRNAs of rice and Arabidopsis (Lu et al. 2015; Ye et al. 2015), maize circRNAs are more likely to be surrounded by retrotransposon LLEs and their reverse complementary pairs, which is similar to what has been observed in humans (Jeck et al. 2013; Zhang et al. 2014) and might be also related to the biogenesis of circRNAs in maize.

### Biogenesis of circRNAs in maize

circRNAs were first reported more than 30 years ago (Hsu et al. 1979). However, besides circular single-stranded RNA genomes of viroids and hepatitis delta virus, and structural circRNAs of tRNAs, rRNAs, which are circularized via ribozymal activity (Group I and Group II introns) (Grabowski et al. 1981; Lehmann and Schmidt 2003), most other endogenous mRNAs were usually thought to be by-products of pre-mRNA processing, and thus were considered to be formed by splicing errors (Capel et al. 1993; Zaphiropoulos. 1993; Pasman et al. 1996). Nevertheless, deep high-throughput sequencing and robust bioinformatic analyzing methods have detected hundreds of thousands of exonic and intronic endogenous circRNAs (Jeck et al. 2013; Zhang et al. 2014; Lu et al. 2015; Ye et al. 2015), which are abundant and even more stable than their parental linear RNAs. Ample evidence has implied that endogenous circRNAs may not solely be formed by splicing error but by certain biological mechanisms.

circRNAs have been reported to be surrounded by long introns, and it has been reported that intronic elements are sufficient to promote circularization in the case of *sex-determining region Y* (*Sry*) circRNA (Dubin et al. 1995). Recently, Jeck and colleagues (2013) showed that introns flanking circRNAs are enriched in ALU repeats in humans. Furthermore, circRNAs are generated cotranscriptionally with linear transcripts and that their production rate is mainly determined by intronic sequences (Ashwal-Fluss et al. 2014). Moreover, Zhang and colleagues (2014) analyzed the enrichment of transposon ALU and their reverse complementary structures, and validated that the complementary sequence could mediate RNA circularization. Finally, lariat-driven circularization and intron pairing-driven circularization were proposed for the biogenesis of circRNAs (Chen et al. 2015; Chen et al. 2016).

Here, we primarily focused on the exonic endogenous circRNAs and bioinformatic analyses of flanking genomic sequences of circRNAs uncovered that retrotransposon LLEs and LLERCPs were significantly enriched around circRNAs, which is concordant with the intron-driven circularization model. Notably, not all the genes with LLERCPs have detectable circRNAs in our study. This may be due to the fact that these genes might not be expressed in our sample given the strong tissue-specific expression feature of circRNAs. Meanwhile, we also noticed that about 80% do not have LLERCPs. These circRNAs may be caused either by reverse complementary sequences of other repetitive elements (Fig S8) or other biological mechanisms. Of particular note, there is also an enrichment of reverse complementary pairs in genes with detectable circRNAs, which are over 10 Kb from circRNA junction sites, which indicates that genomic structure especially the 3-dimension of genome (3-D) structure may play a role in the formation of circRNAs. Since maize lacks 3-D genome data, we analyzed 3-D data and circRNAs in humans. By taking the human GM12878 1kb resolution intra-chromosomal contact matrix (Rao et al. 2014) and human GM12878 circRNAs (CIRCpedia, http://www.picb.ac.cn/rnomics/circpedia/) together, we found that 90.83% circRNA’s splice sites had spatial interaction, suggestive of potential relationship between circRNAs and genome three-dimensional structure. Our results not only provide more evidences that reverse complementary sequences can mediate the cotranscription of circRNAs and linear RNAs, but also highlight the important roles of 5’ and 3’ regions of genes in the formation or regulation of circRNAs and linear transcripts in maize.

### A new functional role in phenotypic variation for transposons in maize

Transposons or transposon-like repetitive elements constitute more than 85% of maize genome (Schnable et al. 2009). These elements dramatically diversified the maize genome, creating or reversing mutations, causing presence/absence variation or copy number variation of genomic regions, and even altering genome size among maize inbreds, which results in the maize genome exhibiting dramatic diversity among inbreds (Lai et al. 2010). Accordingly, maize phenotypic variation could be caused by the insertion of transposons either in the protein coding regions with amino acid alternation, or in flanking regions of functional genes with epigenomic variation, which can further alter nearby gene expression (Wei and Cao 2016). Besides, phenotypic variation could result from the small interfering RNAs, which were generated by the homologous genomic sequences captured and moved by transposons from original functional genes (Lisch 2009). It has been widely reported that the phenotypic variation of maize plant architecture (*tb1* – tillering; Wang et al. 1999), drought resistance (*ZmVPP1;* Wang et al. 2016), pathogen resistance (Yang et al. 2013; Zuo et al. 2015) were associated with transposons and caused by one of the above functional ways of transposons.

In our study, we observed that genes with circRNAs are more likely to be associated with trait-associated sites (TAS), which were identified by whole genome association mapping. Most intriguingly, about 17% these circRNA genes with TAS are associated with LLERCPs, indicating that transposons may function to form circRNAs to titrate functional gene expression and result in increased phenotypic variation in maize. Given the dramatic genomic structural variation among maize inbreds, which is primarily caused by transposons, we propose a new functional role of transposons to modulate phenotypic variation through the formation of circRNAs and regulation of gene expression in maize (Fig 8). For a gene affecting plant height, absence of LLERCP will only produce linear transcript and lead to a certain amount of plant height. When there are LLERCPs, both circRNA and linear transcript could be produced, and the circRNA could interact with the linear transcript either reducing or increasing its expression, resulting in variations in plant height. We validated this proposed model for a particular circRNA, circ352, which is derived from a plant height-associated gene. Taken together, our study uncovered a potential new functional role of transposons to form circRNAs to modulate phenotypic variation in maize, which may also exist in other plants or animals.

**Fig 8.**
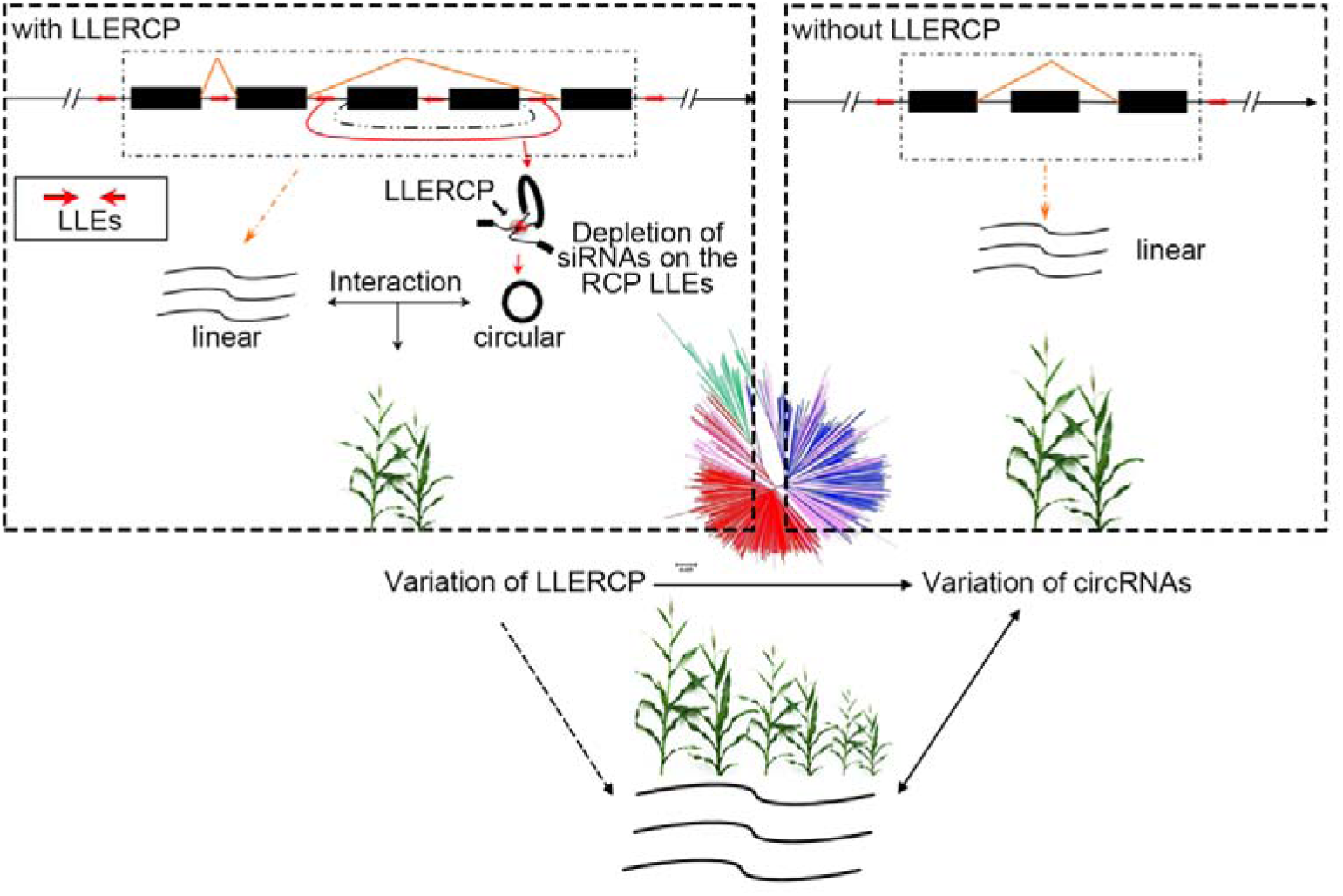
A proposed model where the variation of RCPLLEs among inbreds mediates the interaction of circRNAs and functional linear transcripts and thereby affects phenotypic variation in maize.

## Methods

### Plant materials

Seeds of B73 were planted at room temperature of 24°C in black pots with 16 h light/8h dark in Huazhong Agriculture University, Wuhan, China. The third leaves of V3 stage seedlings were harvested and 2 plants were pooled for RNA isolation.

### Total RNA, genomic DNA extraction, and RNase R treatment

Total RNA was isolated from the pooled sample using TRIzol reagent (Invitrogen, Shanghai, CN) according to the manufacturer’s protocol. To pyrolyze the nucleoprotein, 0.20 ml of trichloromethane was added, followed by 0.20 ml isopropanol to precipitate RNA. And 1ml of 75% ethanol was added to remove the other impurities. Total RNA samples were treated with DNase I (NEBBeijing, CN) and precipitated by sodium acetate-ethanol to remove DNA contamination and salts. RNA was evaluated using 1.0% TAE–agarose gel electrophoresis. For RNase R-treated total RNA samples, the purified DNaseI-treated total RNA was incubated for 15 min at 37°C with 3 units RNase R per μg of total RNA(Epicentre, Shanghai, CN). RNA was subsequently purified by sodium acetate-ethanol method. Total RNA with (RNase R +) or without (RNase R-) RNase R treatment samples were reverse transcribed with random primers using SSIII reverse transcriptase (Invitrogen, Shanghai, CN). Genomic DNA of maize was extracted using the conventional cetyltrimethylammonium bromide (CTAB) method.

### Library preparation and Illumina sequencing

Both RNase R treated RNA (Ribobio, Guangzhou, CN) and poly (A) selected RNA (ANOROAD, Beijing, CN) samples were used for library preparation according to Illumina library construction protocols, followed by circRNA-Seq and mRNA-Seq, respectively. Libraries were sequenced for both ends using the Illumina Hi-Seq3000 with 150 cycles.

### Public datasets used for the identification of circular RNAs

Public maize RNA-seq datasets were obtained from Sequence Read Archive (SRA) database and these include 368 lines of an association panel (kernels; SRP026161), 503 diverse inbred lines (seedings; SRP018753) and various tissues of B73 (leaf, seed, embryo, seeding, endosperm, embryo sac, ovule, pollen, root and shoot apical meristem) (Table S4).

### Computational pipeline of detecting circular RNAs

All three public RNA-seq datasets were first preprocessed using FastQC (http://www.bioinformatics.bbsrc.ac.uk/projects/fastqc/; v0.11.3) and Trimmormatic (v0.33) (Bolger et al. 2014). Briefly, raw reads were trimmed to remove the overrepresented sequences, adaptor contaminations and poor-quality base calls. For the identification of circRNAs, several challenges, including discriminations of trans-splicing, genetic rearrangements, sequencing errors, alignment errors, and *in vitro* artifacts, would result in high false discovery rate (Chen et al. 2015). Two major methods, including pseudo-reference-based strategy and fragment-based strategy, have been devised to detect circRNAs. To circumvent such challenges, different methods were employed for different transcriptome datasets. For each RNA-seq sample, KNIFE (pseudo-reference-based strategy, Szabo L et al. 2015) was used for the genome-wide identification of circRNAs. The pseudo-reference was built with default parameters (the flanking length of each splice site is 150 bp, the maximum distance of any pair of splice sites is 1 Mb). The candidate circRNAs were identified by the naive method (p-value) for this junction based on all aligned reads (p>=0.9) and then merged into one non-redundant set. To reduce the false-discovery rate (e.g, sequencing artifacts), we filtered the circRNAs which appeared less than 10 times in all the above samples or were originated from two genes. Based on paired-end information, we also filtered the candidates which had less than 90% reverse-read supported. Considering that the short length of the exon/UTR circular may be also misleading in scrambled-junction reference database procedure, we the exon/UTR circRNAs, which is less than 50bp. Finally, to alleviate the influence of repetitive sequences, junction reads were re-mapped to B73 reference sequence (v3.26) by using BLAT (Kent 2002). And B73 reference sequences annotated by TRF (Benson 1999) as a tandem repeat were also excluded. For the datasets collected by circRNA-Seq, CIRIexplore2 (fragment-based strategy, Zhang et al. 2016) were employed for the identification of circular RNAs (Fig S1). CircRNAs with at least 2 junction reads, were considered as high-confidence circRNAs in maize.

### RNA-seq analysis

RNA-seq reads were first preprocessed using FastQC (v0.11.3) and Trimmormatic (v0.33) (Bolger et al. 2014), and then aligned to B73 reference sequence (v3.26) using TopHat2 (v2.1.0) (Kim et al. 2013) and transcript abundances were estimated by Cufflinks (v2.2.1) (Trapnell et al. 2010). TPM were calculated by custom scripts.

### Validation of circRNAs

To validate the maize circRNAs we identified, seeds of B73 were germinated at the same condition of samples that have been subjected to circRNA-Seq and mRNA-seq. At the V3 stage the third leaves were harvested. Total RNA was isolated from the pooled sample using similar protocol as well. The PCR primers were designed for the validation of 19 randomly-selected circRNA (Table S2). cDNA (RNase R+), genomic DNA and total RNA were used as templates for the validation of circRNAs by quantitative PCRs. The reagent of 2x Taq Master Mix(Vazyme, Nanjing, CN) was used for cDNA and gDNA amplification with the touchdown PCR program to detect the candidate circRNA templates. Then, Sanger sequencing were performed on all PCR products.

### Characteristics analysis of circular RNAs

The gene length of circular/linear genes was calculated based on the B73 annotations (AGPv3, Schnable et al. 2009). As control, Monte Carlo simulation method was used for linear transcripts. Briefly, 1,245 (same as circular genes) genes without detectable circRNAs were randomly selected and 100 simulations were performed. Then, maximum and minimum average-gene-length of each of the 100 simulations were summarized. Lastly, we repeated the process 1000 times and compared the average gene length of circular RNA to the 1000*100 simulations. Similar method was used for the comparison of other genomic features between circRNAs and linear transcripts.

### MicroRNA target analysis

To compare the genes with and without detectable circRNAs, we randomly selected 1,245 genes without detectable circRNAs, 1,245 genes with LLERCPs, and 1,245 genes with same gene-length distribution as circular genes, accordingly. Furthermore, psRNATarget (Dai and Zhao 2011) were used for microRNA target prediction, 321 putative miRNAs of *Zea mays* were used as the preloaded small RNAs.

### Transposon analysis

To detect the relationship between circular RNAs and the transposons, repetitive information of B73 reference sequence (v3.26) were obtained using RepeatMasker (v2.1; -species maize; http://www.repeatmasker.org/), and custom scripts were used to extract the annotation of repetitive elements in the flanking regions of 5’ and 3’ splicing sites. As a control, Monte Carlo simulation method was performed again for the comparison between genes with and without detectable circRNAs as described before.

### Small RNA analysis

B73 small RNA datasets were download from Gene Expression Omnibus (GEO) database under the accession number GSE39232 (Regulski et al. 2013). Only 21-25 nt small RNAs were used in downstream analysis. Raw reads were first preprocessed using FastQC (v0.11.3) and Trimmormatic (v0.33) (Bolger et al. 2014). The trimmed reads were aligned to B73 reference sequence (v3.26) with Bowtie (v0.12.9; -n 0) (Langmead et al. 2009). And the small RNAs mapped on LLEs were extracted and summarized using SAMtools (v0.1.19) (Li et al. 2009) and the normalized expression-level of LLEs were calculated as:

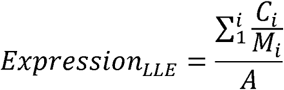

(i, the number of distinct small RNA reads aligned to specific LLE; C_i_, the abundance of a specific distinct small RNA; M_i_, the number of genomic loci that the distinct small RNA read has been mapped; A, total number of mapped small RNAs).

### Discovery of potential functional roles for circRNAs

To test if circRNAs play a role in phenotypic variation, genes with trait-associated sites were collected from four different genome-wide association mapping (GWAS) studies (Li et al. 2013; Yang et al. 2014; Wen et al. 2014; Wallace et al. 2014). Genes with significantly trait associated sites are more likely to be associated with phenotypic variation. Thus, trait-associated genes could be a good resource for the functional enrichment analysis of circRNA genes. Similarly, Monte Carlo simulation method was employed for the comparison of appearance frequency in the trait-associated gene list between linear and circular genes. To rule out the spurious enrichment of circRNA genes in the trait-associated gene list due to the longer gene length of circRNA genes, we performed the functional enrichment analysis for genes with detectable circRNAs, randomly-selected genes with same gene-length distribution as circRNA genes, randomly-selected genes with LLERCPs but without detectable circRNAs, and randomly-selected genes at the same time. Additionally, Gene ontology (GO) enrichment analyses of circular gene were performed using agriGO (Du et al. 2010) with singular enrichment analysis and whole set of the annotated genes as the reference.

### Association analysis between presence/absence of LLERCPs and plant height variation in maize

Previous study has identified a functional gene *GRMZM2G089149,* that was associated with plant height variation in a US association panel (Wallace et al. 2013). Here, we found that a cirRNA-circ352 is derived from this plant-height associated gene and has intact LLERCPs around. To test the relationship between LLERCPs and phenotypic variation, we randomly selected 38 diverse inbreds from a Chinese association panel (Yang et al. 2014), designed primers to amplify the intact LLERCPs of circ352, and quantified the relative expression levels of circ352 across different inbreds in 3^rd^ leaf of seedlings. The presence/absence of LLERCPs across 38 diverse inbreds was detected using PCR amplification and Sanger sequencing. T-test was employed to test the plant height difference between inbreds with LLERCPs and without LLERCPs.

## Data access

The datasets of CircRNA-Seq (PRJNA356366) and poly (A) selected mRNA-Seq (PRJNA356498) were generated and deposited to Sequence Read Archive database (https://www.ncbi.nlm.nih.gov/sra).

## Acknowledgments

This research was supported by the National Key Research and Development Program of China to Mingqiu Dai (2016YFD0100600), and the National Key Research Project of China – Molecular Mechanisms conferring heterosis in staple crops to Lin Li (2016YFD0100802). This research was also supported by Huazhong Agricultural University Scientific & Technological Self-innovation Foundation to Lin Li (Program No. 2015RC016).

## Author contributions

L. L. and M. D. designed and supervised this study. L. C., P. Z., Y. F. and Q. Lu performed the data analysis and circular RNA validation. J. H. performed RNA extraction and purification. L. C. and L. L. wrote the manuscript. P. S. S., G. J. M., Q. Li and J. Y. reviewed and edited the manuscript.

